# MicroGenomer: A Foundation Model for Transferable Microbial Genome Representations Enabling Multi-scale Genomic Understanding and Ecophysiological Trait Prediction

**DOI:** 10.64898/2025.12.28.696777

**Authors:** Qiang Kang, Yucheng Guo, Bozhen Hu, Xiaoping Li, Qirong Yang, Yujie Yang, Dongli Lu, Haotong Sun, Ruilai Xu, Ziyao Xu, Haochen Wang, Yuheng Lin, Yuxiang Li, Yong Zhang, Feiran Li, Xiaoming Zhang, Peng Yin

**Affiliations:** BGI Research, Wuhan, Hubei, China, 430074; BGI Research, Shenzhen, Guangdong, China, 518003; BioMap Research, Beijing, China, 100086; Laboratory of Integrative Biomedicine, Department of Biology, University of Copenhagen, Copenhagen, Denmark; Shenzhen Key Laboratory of Environmental Microbial Genomics and Application, BGI Research, Shenzhen, Guangdong, China, 518003; School of Artificial Intelligence, University of Chinese Academy of Sciences, Beijing, China, 100049; College of Wildlife and Protected Area, Northeast Forestry University, Harbin, Heilongjiang, China, 150040; Institute of Biopharmaceutical and Health Engineering, Tsinghua Shenzhen International Graduate School, Tsinghua University, Shenzhen, Guangdong, China, 518055

**Keywords:** Foundation model, Microbial genome representation, Multi-scale genomic understanding, Ecophysiological trait prediction

## Abstract

Microorganisms underpin global biogeochemical cycles and represent a vast, underexplored reservoir for sustainable biotechnology and human health. Their diversity, arising from the interplay of numerous genes within holistic genomic contexts, dictates complex ecophysiological traits across varied environments. To bridge the gap between complex genomic sequences and associated biological functions and phenotypic outcomes, we present MicroGenomer, a foundation model for transferable microbial genome representations enabling multi-scale genomic understanding and ecophysiological trait prediction. MicroGenomer leverages a hierarchical training strategy comprising pre-training on large-scale genomic sequences (234.5 billion base pairs), domain-specific mid-training using the GTDB-curated marker gene set, and task-specific post-training. Despite a streamlined architecture of 470 million parameters, MicroGenomer achieves performance on key tasks competitive with models nearly 85 times its size. Advancing beyond established gene-scale encoders, MicroGenomer generates robust embeddings at the genome scale for downstream modeling. Extensive evaluations demonstrate that MicroGenomer effectively captures phylogenetic structures in species space, excelling in gene- and genome-scale understanding as well as ecophysiological trait prediction. The practical utility of these capabilities is further demonstrated by targeted wet-lab validation on newly isolated strains, indicating that MicroGenomer’s predictions provide reliable guidance for biological experiments. Collectively, MicroGenomer offers a high-performance, resource-efficient framework that transforms raw sequence data into actionable biological insights, providing a powerful foundation for microbiome research and biotechnology.

## 1 Introduction

Microorganisms, as fundamental components of Earth’s ecosystems, drive global biogeochemical cycles and profoundly influence human health, agricultural productivity, and environmental sustainability [1]. High-throughput sequencing and genome assembly have generated vast amounts of genomic and metagenomic data across diverse environments, including oceans, soils, and the human gut [2, 3]. Large-scale resources such as the Genome Taxonomy Database (GTDB) [4] and MGnify [5] have enabled systematic phylogenetic reconstruction and functional annotation, revealing the immense functional potential encoded within complex genomic contexts. However, bridging the gap between raw sequences and a systemic understanding of microbial life remains a significant challenge. Establishing the mechanistic links that span from gene scale to genome scale is essential for understanding how specific taxa manifest complex ecophysiological traits [6]. Microbial technologies already play a pivotal role in industrial fermentation and probiotic discovery [7], carrying immense economic weight (e.g., the global probiotics market exceeded US$60.5 billion in 2022, and China’s biofertilizer sector alone comprises over 10,000 registered products [8]). These statistics highlight the growing bioeconomic impact of microbial applications and the necessity for precision modeling. The translation of these genomic resources into actionable insights often involves computational frameworks designed to predict complex traits from sequence data. Yet, many existing methods rely on engineered sequence features and supervised models trained on limited, imbalanced labels, which can lead to poor generalization across diverse species or environments [9]. This emphasizes the urgent need for unified and transferable representations that can integrate information from individual genes to entire genomes, enabling robust multi-scale analysis and ecophysiological trait prediction across varied taxa and habitats.

Artificial intelligence (AI) foundation models have demonstrated broad utility across various biological domains, facilitating discoveries from protein structure prediction to complex microbial community analysis. At the protein level, models such as AlphaFold [10], RoseTTAFold [11], and evolutionary-scale language models like ESM-2 [12] have shown that large-scale pre-training can extract rich structural and functional representations from sequence data. At the cellular level, frameworks like scGPT have successfully generalized across diverse single-cell omics tasks [13]. Paralleling these developments in genomics, DNA language models such as DNABERT [14], Nucleotide Transformer (NT) [15], and HyenaDNA [16] treat DNA sequences as a structured language, capturing long-range regulatory patterns and k-mer signatures. More recently, large-scale generative models and those trained on metagenomic scaffolds have begun to decode microbial co-regulation and functional relationships [17]. These collective advances suggest that self-supervised learning can yield highly reusable sequence embeddings, significantly reducing the historical dependence on large, manually labeled datasets.

Despite this progress, current DNA language models remain limited in microbial contexts, especially when moving from sequence patterns to functional understanding. Most existing frameworks, including long-context models such as EVO and Hye-naDNA, primarily operate as sequence-level encoders that treat genomic data as continuous strings of nucleotides. While these models are effective at capturing structural motifs within their window constraints, they lack an explicit mechanism to prioritize and integrate discrete coding sequences (CDSs), which serve as the primary drivers of microbial function. This gene-agnostic approach often fails to bridge the gap between local genetic variation and the collective functional potential of a complete genome. Furthermore, microbial genomes exhibit a highly organized hierarchical structure where operons and gene clusters cooperate to dictate systemic traits. Few models are designed to explicitly aggregate these gene-scale signals into unified representations that encompass the entire genomic scale. The persistent sparsity and taxonomic bias of microbial phenotype labels also make traditional supervised training on raw sequences prone to overfitting. While specialized tools like Traitar or gRodon demonstrate the potential for trait inference, they are often restricted to narrow tasks and lack the transferability required for diverse ecological applications [18, 19]. These cumulative challenges necessitate a framework that transcends simple sequence encoding to achieve a multi-scale and phylogeny-aware understanding of microbial genomes.

To address these limitations, we present MicroGenomer, a foundation model for transferable microbial genome representations that enables multi-scale genomic understanding and ecophysiological trait prediction. MicroGenomer is built upon a Transformer-based architecture and follows a three-stage training pipeline leveraging extensive genomic corpora totaling hundreds of billions of base pairs (bp). Following large-scale pre-training on raw sequences, the model undergoes a domain-specific midtraining stage using marker gene sets to bridge the gap between sequence and function, followed by a task-specific post-training stage for specialized downstream applications. Crucially, MicroGenomer transcends traditional gene-centric approaches by uniquely aggregating functional embeddings of all constituent CDSs into robust, genome-scale representations. With approximately 0.47 billion parameters, MicroGenomer achieves performance competitive with the EVO series models nearly 40 billion in size across some key benchmarks. We evaluate the model’s capabilities across a broad spectrum of tasks, ranging from gene-scale mutational effect prediction to genome-scale metabolic model analysis and ecophysiological trait prediction (such as optimal growth conditions). Furthermore, the practical utility of MicroGenomer is validated through wet-lab cultivation of newly isolated strains, where its predictions provided effective guidance for determining optimal growth pH and temperature. By transforming raw sequence data into actionable biological insights, MicroGenomer provides a resource-efficient foundation for microbiome research, strain engineering, and microbial biotechnology.

## 2 Results

### 2.1 Overview of MicroGenomer

MicroGenomer is a domain-adaptive transfer learning framework with 470 million parameters designed to generate transferable microbial genome embeddings ebabling multi-scale genomic understanding and ecophysiological trait Prediction. It is trained in three successive stages: pre-training, mid-training, and post-training.

In the pre-training stage, DNA sequences from the OpenGenome dataset (234.5 billion nucleotides) are tokenized at single-nucleotide resolution, yielding approximately 1,080 billion tokens. We then train a 24-layer Transformer encoder, serving as the MicroGenomer core, on this corpus with a context window of 8,192 tokens using a masked language modeling (MLM) loss (ℒ_MLM_). The resulting pre-trained model generates high-resolution gene-scale embeddings, enabling the execution of tasks such as mutational effect prediction and the Genome Understanding Evaluation (GUE) benchmarks.

In the mid-training stage, we use a curated GTDB microbial marker gene set (36 billion nucleotides) comprising CDSs from approximately 110k genomes across 53 archaeal and 120 bacterial marker gene families. These CDSs are tokenized at singlenucleotide resolution, and the pre-trained MicroGenomer core is further optimized on this corpus with a maximum context window of 2,000 tokens. For each genome, the generated CDS embeddings (gene-scale) are aggregated via pooling into a unified genome-scale embedding. This stage is optimized using a phylogeny-based representation distance loss (ℒ_species_) and a gene-category classification loss (ℒ_CLS_). The resulting genome-scale representations support tasks including metabolic model analysis and the cluster alignment of embeddings vs. phylogeny.

**Fig. 1:**
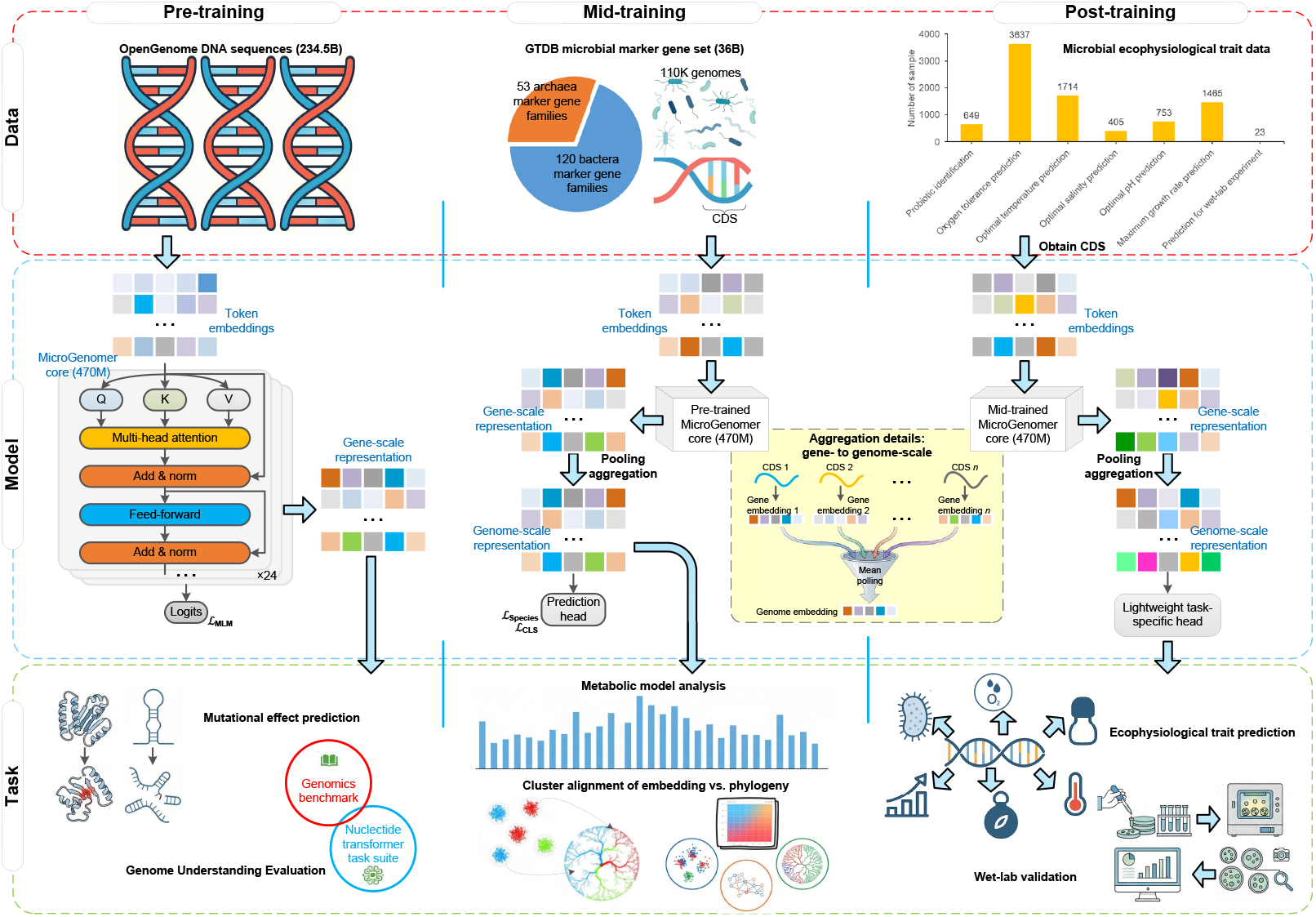
Overview of the MicroGenomer framework. The model follows a three-stage training strategy to generate hierarchical embeddings spanning from gene-to genome-scales. MicroGenomer supports diverse evaluations, ranging from gene-scale tasks (e.g., mutational effect prediction and genome understanding evaluation) to genome-scale applications (e.g., metabolic model analysis and ecophysiological trait prediction.

In the post-training stage, the mid-trained MicroGenomer core serves as a feature extractor to generate fixed genome-scale embeddings for microbial datasets with phenotype annotations. These embeddings serve as inputs for lightweight, task-specific prediction heads, enabling MicroGenomer to complete diverse ecophysiological trait prediction tasks, such as probiotic identity, oxygen tolerance, and optimal growth conditions (temperature, pH, salinity, and maximum growth rate). To ensure robust generalization across clades, these predictions are performed under phylogenetically informed evaluation settings. Finally, MicroGenomer’s predictive power facilitates wetlab validation on newly collected isolates to characterize their environmental growth limits.

### 2.2 MicroGenomer exhibits robust generalization and efficient multi-scale genomic understanding

#### MicroGenomer achieves remarkable zero-shot mutational effect prediction

In biological sequence modeling, learning sequence probabilities from large evolutionary datasets allows models to link mutational effects to biological function [20]. Deep mutational scanning (DMS) provides a systematic framework for quantifying the fitness impact of mutations across diverse proteins and noncoding RNAs (ncRNAs), and is widely used to evaluate both protein sequence models [21, 22] and genomic language models [16, 23, 24]. We compare the pre-trained MicroGenomer core against state-of-the-art models, including the EVO series [24] and MambaDNA [25], specifically assessing their ability to predict mutational effects on prokaryotic proteins and ncRNAs.

As shown in Fig. 2A, MicroGenomer achieves the highest mean Spearman correlation among all tested models in ncRNA mutational effect prediction. Notably, in protein mutational effect prediction, MicroGenomer delivers performance comparable to the top-tier EVO2-40B and surpasses all other baselines. Remarkably, MicroGenomer achieves these results with only 0.47B parameters, a mere 1.18% of the parameter scale of EVO2-40B. These findings demonstrate that MicroGenomer effectively captures the constraints between genetic alterations and functional outcomes through sequence-likelihood inference. Furthermore, this high level of predictive accuracy within a significantly smaller parameter footprint underscores that MicroGenomer offers a highly efficient and robust framework for characterizing mutational impacts across diverse genomic contexts.

**Fig. 2:**
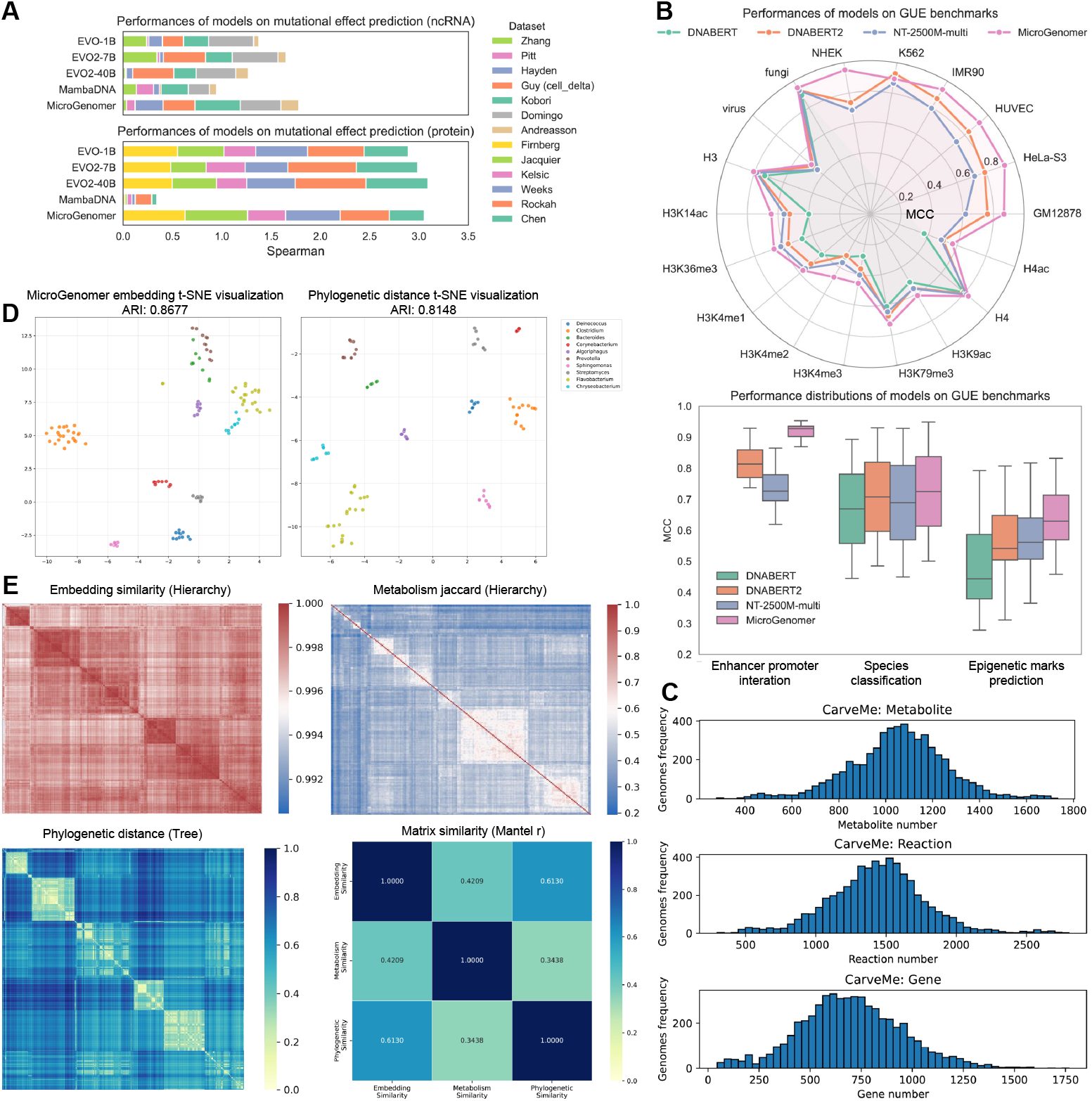
Multi-scale evaluation of MicroGenomer representations across genomic and evolutionary tasks. **(A)** Performance comparison of MicroGenomer and baseline models for zero-shot mutational effect prediction. **(B)** Performance comparison of MicroGenomer and baseline models on GUE benchmarks. **(C)** Distributions of metabolite, reaction, and gene counts across 5,587 bacterial genome-scale metabolic models. **(D)** t-SNE visualizations of representations for genomes from the ten most abundant genera. **(E)** Similarity structure across genomes: cosine similarity of MicroGenomer embeddings, Jaccard similarity of metabolic feature sets, phylogenetic distances from the bac120 tree, and Mantel *r* values between the three matrices (*p* = 0.001).

#### MicroGenomer excels in functional characterization on GUE benchmarks

We evaluate the pre-trained MicroGenomer core on GUE benchmarks, which integrates diverse tasks from the Genomics Benchmark [26] and the NT task suite [27]. To adapt the model to downstream tasks, we employ standard Low-Rank Adaptation (LoRA) fine-tuning [28], a widely used parameter-efficient method. For baseline comparisons, we use performance results reported in the original studies of competing models. We compare MicroGenomer against three prominent baselines, i.e., DNABERT [29], DNABERT2 [26], and NT variants (NT-2500M-multi, 2500M parameters, trained on multi-species data) [27].

As illustrated in Fig. 2B, in the enhancer–promoter interaction prediction across different cell lines (GM12878, HeLa-S3, HUVEC, IMR90, K562, and NHEK), MicroGenomer consistently achieves higher Matthews Correlation Coefficient (MCC) scores than DNABERT2 and NT-2500M-multi, while the original report of DNABERT did not provide results for this task. In the species classification tasks on fungi and viruses, MicroGenomer still achieves higher MCC scores than any other model. For epigenetic marks prediction, MicroGenomer continues to maintain the highest MCC across various histone modifications (H3, H3K14ac, H3K36me3, H3K4me1, H3K4me2, H3K4me3, H3K79me3, H3K9ac, H4 and H4ac). Overall, MicroGenomer demonstrates state-of-the-art performance across multiple genomic tasks in the GUE benchmarks, surpassing other leading models in the field. These results indicate that MicroGenomer not only leverages evolutionary signals during pretraining to handle long sequences but also excels in capturing fine-scale regulatory patterns and annotations.

#### MicroGenomer demonstrates superior precision in capturing evolutionary distances

Beyond sequence-level understanding, MicroGenomer is captures systemic biological patterns, specifically the intricate relationships between phylogenetic evolution and metabolic functions. To evaluate the quality of the genome-scale representations generated by the mid-trained MicroGenomer core, we collect a dataset of 5,587 genome-scale metabolic models for bacterial species. These models, constructed via CarveMe [30] from NCBI RefSeq (release 84) [31], capture metabolic diversity at the strain level. Each metabolic model encompasses information on reactions (with reaction IDs), metabolites (with metabolite IDs referencing species), and genes (with gene IDs combining geneProductRef and geneProduct). Fig. 2C illustrates the distribution of metabolites, reactions, and genes across the dataset. Notably, these distributions are all unimodal (i.e., having a single peak), indicating a concentrated distribution of values. We focus on the top ten genera with the highest number of genomes in this dataset. We conduct t-SNE visualization (Fig. 2D) to compare genome-scale embeddings generated by MicroGenomer with phylogenetic distances [32, 33], and evaluate the clustering performance using the Adjusted Rand Index (ARI). Since the mid-training phase of MicroGenomer incorporates evolutionary distance loss derived from the species phylogenetic tree, the generated embeddings produce clustering patterns that closely mirror the phylogenetic relationships. Moreover, the ARI obtained from clustering based on MicroGenomer embeddings is higher than that obtained from clustering based directly on phylogenetic distances, demonstrating that MicroGenomer not only preserves but also sharpens the evolutionary structure encoded by the phylogenetic tree.

The core objective of hierarchical clustering is to “order” the samples such that similar samples are placed closer to each other after sorting, thereby making subsequent visualizations, such as heatmaps, more intuitive in presenting the clustering patterns among the samples. Fig. 2E (upper left) illustrates the hierarchical clustering results for species similarities, calculated from the cosine similarity among the embeddings of different bacterial genomes generated by the mid-trained model of MicroGenomer. Red hues (approaching 1.000) signify a very high degree of similarity between the corresponding pairs of items, whereas white hues (approaching 0.995) denote relatively lower similarity. The structure of the heatmap reflects the outcome of hierarchical clustering. Within the heatmap, contiguous red-colored blocks represent groups of items that exhibit a high level of mutual similarity, a result stemming from the clustering process. The panels in Fig. 2E use the same sorted indices for a fair comparison. The Jaccard similarity heatmap in Fig. 2E (upper right) measures the similarity between sets of metabolic features across bacterial genome-scale models. Red tones (near 1.0) indicate a very high Jaccard similarity, implying a significant overlap in the metabolic features between the corresponding pairs of models, while blue tones (near 0.2) indicate low similarity, meaning the sets of metabolic features have only a slight overlap. The contiguous red blocks suggest groups of models with relatively high metabolic similarity. In Fig. 2E (lower left), phylogenetic distance is employed to quantify the evolutionary relatedness between organisms; smaller distances imply closer evolutionary relationships, while larger distances imply greater evolutionary divergence. Yellow tones (near 0.0) indicate very small phylogenetic distances, while dark blue tones (near 1.0) indicate large phylogenetic distances. Blocks in this panel, particularly the yellow-colored ones, suggest groups of genomes with relatively small phylogenetic distances (i.e., closely related groups). The phylogenetic distances are derived from the bac120 tree [34]. Fig. 2E (lower right) visualizes the pairwise similarity (measured by the Mantel test [35]) among three types of similarity matrices: the embedding similarity matrix, the metabolic similarity matrix, and the phylogenetic distance matrix. Mantel *r* quantifies the correlation between two matrices; values closer to 1 indicate a stronger positive correlation, while values closer to 0 indicate a weaker correlation. The relatively higher Mantel *r* between the embedding similarity and metabolic similarity (0.4209), compared to that between metabolic similarity and phylogenetic similarity (0.3438), suggests that our proposed model, MicroGenomer, is capable of generating embeddings that are more closely aligned with metabolic characteristics than with the phylogenetic tree.

### 2.3 MicroGenomer enables precise prediction of microbial ecophysiological traits

We evaluate MicroGenomer across six ecophysiological trait prediction tasks derived from three published benchmark datasets, covering both classification and regression in microbial ecology, physiology, and evolutionary biology. The iProbiotics dataset [36] supports probiotic identification as a binary classification task. The GenomeSPOT study [37] provides four prediction tasks, i.e., oxygen tolerance (classification) and optimal temperature, salinity, and pH preference (regression). The sixth task, maximum growth rate prediction, is evaluated using the dataset introduced in Phydon [38], which integrates codon usage statistics with phylogenetic information to improve growth rate estimation. As shown in Fig. 3A, each task has a relatively modest number of labeled genomes, with sample sizes ranging from 405 to 3,637, reflecting the limited availability of experimentally characterized microbial phenotypes. These tasks provide a broad and challenging assessment of whether MicroGenomer can support accurate trait prediction across diverse metabolic, ecological, and evolutionary traits.

**Fig. 3:**
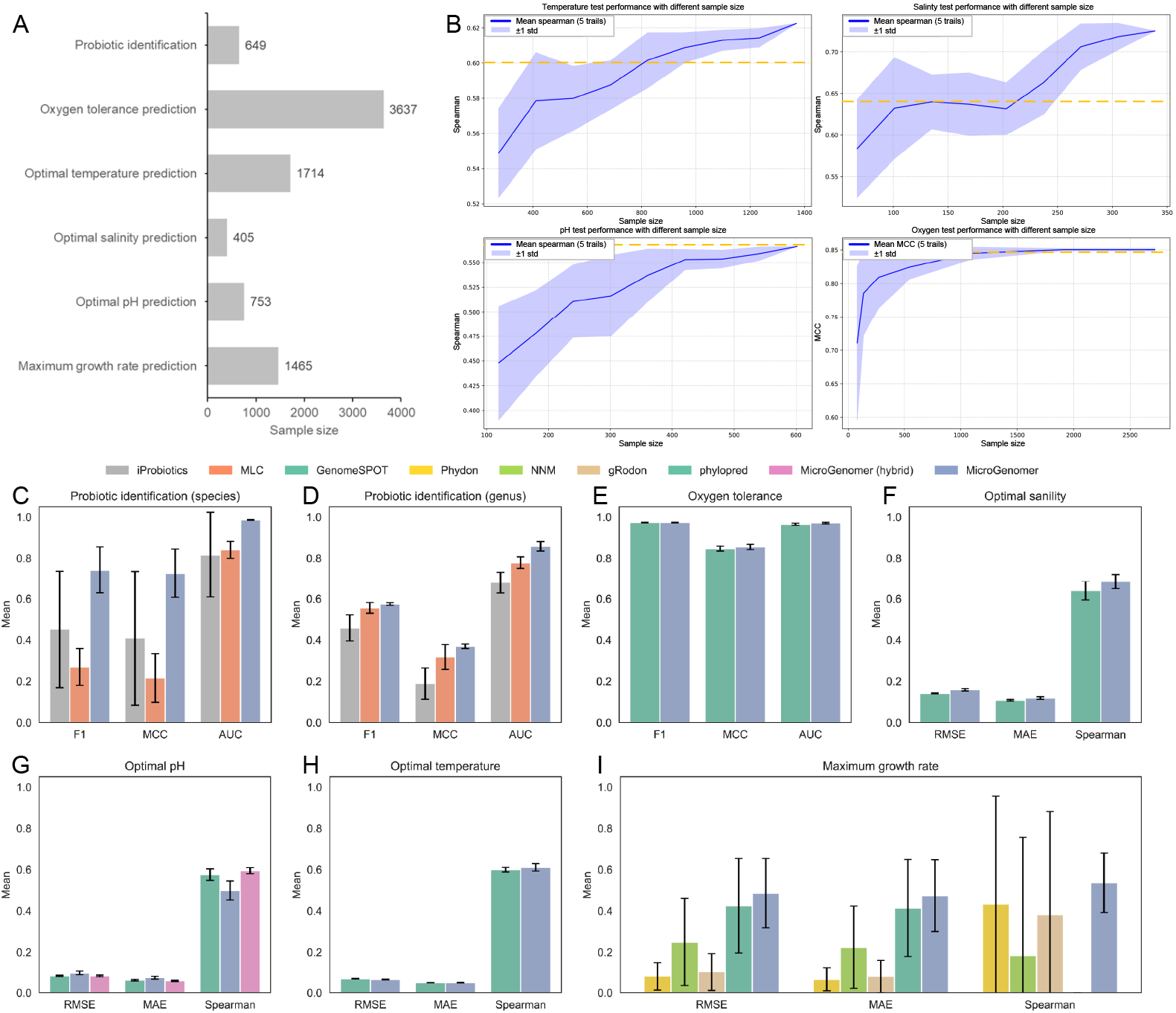
Evaluation of MicroGenomer on microbial ecophysiological trait prediction tasks. **(A)** Sample sizes for six ecophysiological trait prediction tasks. **(B)** Learning curves for MicroGenomer on four trait prediction tasks, showing test performance as a function of the number of training genomes. Solid lines indicate the mean over five independent trials, and shaded areas denote ± 1 standard deviation. **(C)**–**(I)** Comparison of MicroGenomer with baseline methods for probiotic identification at the **(C)** species level and **(D)** genus level, and for the prediction of **(E)** oxygen tolerance, **(F)** optimal salinity, **(G)** optimal pH, **(H)** optimal temperature, and **(I)** maximum growth rate.

For all tasks, we adopt phylogenetically informed data partitioning to ensure rigorous evaluation of MicroGenomer’s generalization ability. For the iProbiotics dataset [36], we first perform a phylogenetically informed split to obtain distinct training and test sets. The training set is then randomly divided into five equal subsets. In each fold, four subsets are used for training, while evaluation is always carried out on the same phylogenetically separated test set. This procedure resembles standard five-fold cross-validation while preserving evolutionary separation between training and test data. Experiments are conducted at both the genus and species levels, and results are compared against iProbiotics [36] and MLC [39]. For the four GenomeSPOT tasks, we adopt the same phylogenetically informed splitting strategy but perform experiments only at the species level, and GenomeSPOT serves as the primary baseline for comparison [37]. For maximum growth rate prediction, we follow the phylogenetically blocked ten-fold cross-validation protocol introduced in Phydon, directly using the fold assignments from the original study and comparing MicroGenomer with Phydon, NNM, and gRodon [38]. This preserves consistency with previous work and enables a fair evaluation of MicroGenomer’s ability to capture growth rate–associated genomic signals.

Given the limited size of these labeled datasets, we do not apply LoRA [28] or other end-to-end fine-tuning strategies to MicroGenomer. Instead, MicroGenomer serves as a fixed genome representation module. For each genome, CDS-derived embeddings are pooled into a single fixed-dimensional vector, which is then used as input to lightweight prediction heads. This design reduces the risk of overfitting, simplifies model comparison across tasks, and cleanly decouples representation learning from downstream inference.

Based on these genome embeddings, we evaluate a diverse suite of machine-learning heads tailored to the task type. For regression-based traits, we consider a two-layer multilayer perceptron (MLP) [40] with a latent dimension of 128, Lasso regression [41], Ridge regression [42], Support Vector Regression (SVR) [43], Random Forest regression [44], k-Nearest Neighbors (KNN) regression [45], and XGBoost regression [46]. For classification tasks, we analogously evaluate a two-layer MLP, logistic regression, Random Forest classification, Support Vector Machine (SVM) classification [47], KNN classification, and XGBoost classification. Hyperparameters for all heads are tuned via grid search [48] on the training/validation splits, and for each method–task pair we report the performance of the best-performing head. The same protocol is applied to MicroGenomer embeddings and to baseline feature representations, ensuring a fair comparison of downstream modeling capacity.

#### Effect of training sample size on MicroGenomer’s performance

In the GenomeSPOT setting, the labeled datasets for each trait contain only hundreds to a few thousand genomes. For deep learning models, limited training sets are highly susceptible to both underfitting and overfitting, which can compromise the reliability of the results. Consequently, we investigate how the training sample size affects MicroGenomer’s performance on the test set, as shown in Fig. 3B.

For optimal temperature prediction (Fig. 3B, upper left), as the number of training genomes increases from roughly 400 to 1,400, the Spearman correlation steadily rises and the blue shaded region ( ± 1 standard deviation over five independent trials) becomes narrower, indicating more stable performance. MicroGenomer’s performance surpasses that of GenomeSPOT once the training set size exceeds around 800 genomes. In optimal salinity prediction (Fig. 3B, upper right), when the number of training genomes increases from about 50 to 350, the Spearman correlation improves from ∼ 0.59 to above 0.70, and MicroGenomer overtakes GenomeSPOT at intermediate sample sizes. For optimal pH prediction (Fig. 3B, lower left), as the training set grows from about 100 to 600 genomes, the Spearman correlation increases from ∼ 0.45 to about 0.57, and MicroGenomer approaches the performance of GenomeSPOT. For oxygen tolerance prediction (Fig. 3B, lower right), increasing the training set from a few hundred to around 2,500 genomes raises the MCC from ∼ 0.71 to about 0.85, with the shaded region again narrowing. At larger sample sizes, MicroGenomer’s MCC matches or slightly surpasses that of GenomeSPOT.

Overall, across all four tasks, increasing the number of training genomes improves MicroGenomer’s performance and reduces performance variability. For optimal temperature, salinity, and oxygen tolerance, MicroGenomer meets or exceeds GenomeSPOT once sufficient training data are available, while for optimal pH it narrows the performance gap as data increase. This highlights the importance of data volume for reliable microbial trait prediction under phylogenetically informed evaluation.

#### Results of different methods on probiotic recognition

We compare MicroGenomer with iProbiotics [36] and MLC [39] on probiotic identification at the species and genus levels. For each method, we report the mean and standard deviation over five independent runs for three metrics: F1-score, MCC, and AUC. At the species level (Fig. 3C), MicroGenomer achieves the best overall performance across all three metrics, substantially outperforming both iProbiotics and MLC, indicating that its genome representations provide a more reliable decision boundary for identifying probiotic strains. At the genus level (Fig. 3D), as expected from the increased complexity of finer-grained genomic variations across diverse genera, all methods experience a decrease in performance. Even under this stricter setting, MicroGenomer remains the most robust method overall. The consistent advantage of MicroGenomer across both taxonomic levels demonstrates the significant benefits of leveraging high-dimensional genome embeddings for ecophysiological trait classification.

#### Results of different methods on oxygen tolerance, optimal salinity, pH, and temperature prediction

We evaluate MicroGenomer on four key ecophysiological traits, oxygen tolerance, optimal salinity, pH, and temperature, benchmarking it against GenomeSPOT, a specialized feature-based approach.

For oxygen tolerance prediction (Fig. 3E), MicroGenomer and GenomeSPOT achieve similar F1-scores, but MicroGenomer attains higher MCC and AUC. This indicates that, at comparable overall sensitivity, MicroGenomer yields better-calibrated decision boundaries and fewer systematic misclassifications between aerobic and anaerobic lifestyles.

For optimal salinity prediction (Fig. 3F), although GenomeSPOT obtains lower Root Mean Squared Error (RMSE) and Mean Absolute Error (MAE), suggesting an advantage in predicting absolute normalized values, MicroGenomer reaches a higher Spearman correlation with the true labels (0.685 vs. 0.640). Given that the primary objective is often to rank strains by their salinity preference rather than to recover exact numeric values, this superior Spearman correlation demonstrates that MicroGenomer provides a more faithful ordering of salinity tolerances across genomes.

The task of optimal pH prediction (Fig. 3G) presents a unique challenge due to the limited number of labeled genomes, where the genome-only MicroGenomer initially lags behind GenomeSPOT. This gap likely stems from GenomeSPOT’s use of specialized subcellular-localization–aware features, which contrast amino acid compositions between intracellular, extracellular, and membrane compartments to capture pH-sensitive adaptation signals. To leverage this complementary information, we construct a hybrid MicroGenomer by augmenting genome embeddings with these protein-derived descriptors (e.g., sequence composition and compartment-specific features). This hybrid variant effectively closes the gap and performs comparably to or slightly better than GenomeSPOT, highlighting the strong compatibility between MicroGenomer’s learned representations and engineered biological features.

For optimal temperature prediction (Fig. 3H), MicroGenomer obtains lower RMSE and higher Spearman correlation, while the MAE results are comparable between the two methods. This performance indicates that MicroGenomer effectively captures thermal adaptation signals directly from genomic sequences and generalizes robustly under phylogenetically informed evaluation.

Taken together, these results show that although protein-level features remain useful for certain traits, particularly under severe data limitations, genome-level embeddings from MicroGenomer already provide competitive or superior performance on most GenomeSPOT tasks, and can be further strengthened by incorporating protein features when appropriate.

#### Results of different methods on maximum growth rate prediction

For maximum growth rate prediction (Fig. 3I), MicroGenomer achieves the highest Spearman correlation among all tested methods, indicating its robust ability to rank microbial strains by their growth potential. While Phydon performs better in terms of RMSE and MAE, reflecting a more precise calibration of absolute growth values, MicroGenomer’s superior rank-based performance is particularly significant. In many ecological and evolutionary contexts, the primary objective is to correctly order strains from slow to fast growers rather than to recover exact numerical rates. Thus, MicroGenomer excels in capturing the genomic signals associated with growth-rate variation, though its absolute accuracy can potentially be further refined by integrating task-specific codon usage or phylogenetic features.

### 2.4 MicroGenomer reveals interpretable patterns of metabolic similarity

For the metabolic similarity prediction, we utilize a metabolic similarity dataset comprising 5,587 bacterial genome metabolic models (Fig. 2C). While MicroGenomer typically aggregates gene embeddings into a genome representation via mean pooling, we implement an attention pooling module to enhance model interpretability (Fig. 4A), This module assigns a learnable weight to each gene while maintaining the original architecture’s integrity. For each genome, the sequence of gene embeddings (dimension *D*) passes through a two-layer MLP (*D* → 64 → 1) to generate scalar attention scores. Following masking and softmax normalization across all gene positions, these weights produce a weighted average of gene embeddings, resulting in a fixed-dimensional aggregated genome representation. This representation is then fed into a two-layer prediction head (*D* → 128 → 1) to output the final metabolic similarity score.

**Fig. 4:**
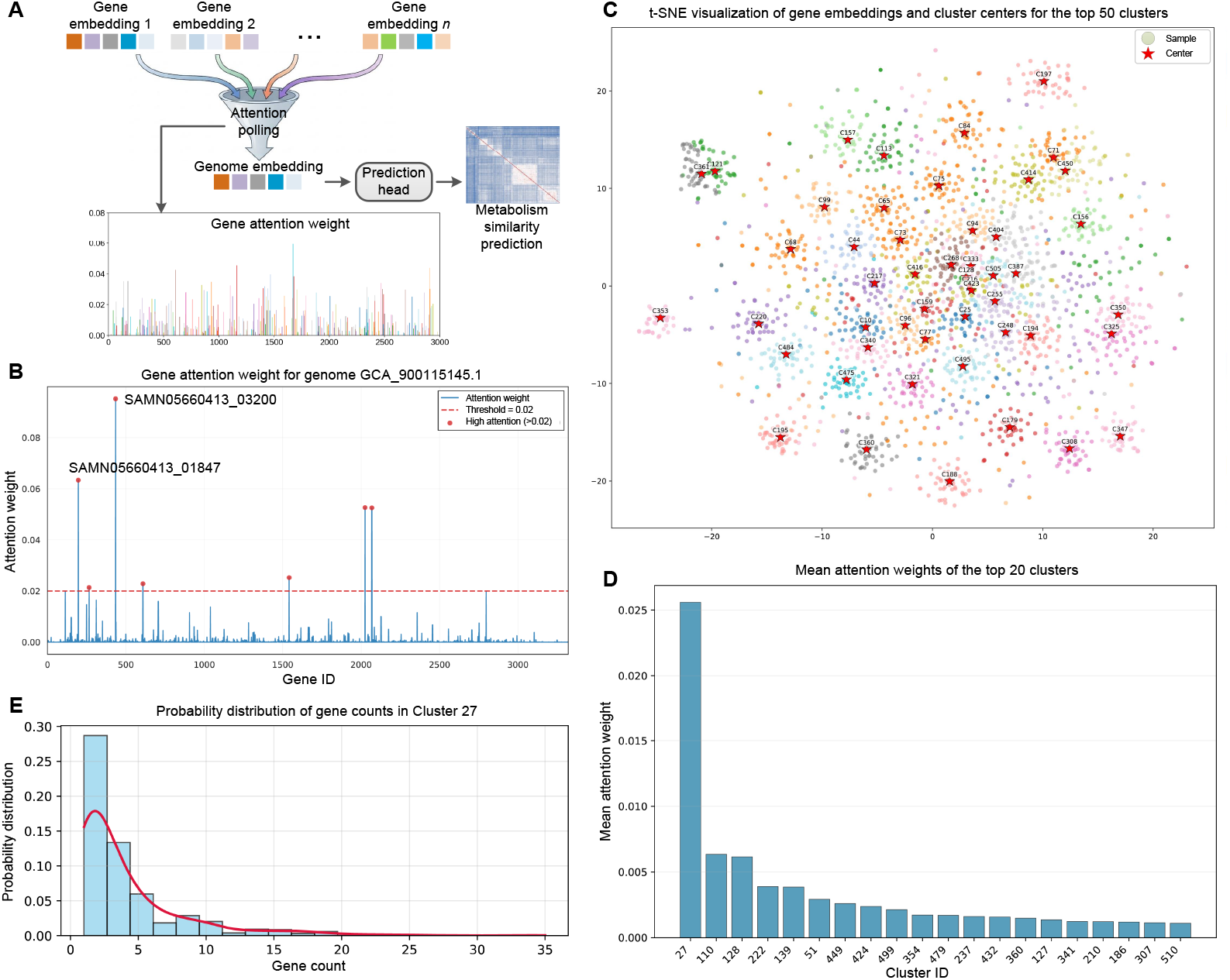
Interpretability analysis of MicroGenomer representations on the metabolic similarity prediction. **(A)** Schematic of the attention pooling module designed to aggregate gene-scale representations into genome-sacle representations. **(B)** Gene attention weights for a representative genome (GCA 900115145.1), where the dashed line indicates the threshold for identifying high-attention genes. **(C)** t-SNE visualization of gene embeddings and cluster centers for the top 50 clusters ranked by the number of member genes. **(D)** Ranked mean attention weights for the top 20 gene clusters, highlighting Cluster 27 as a dominant functional group that drives genome representations. **(E)** Probability distribution of gene counts within Cluster 27 across 898 representative metabolic models.

To visualize the model’s decision-making process at the genome scale, we examine the gene attention weights for a representative genome, GCA 900115145.1 (Fig. 4B). The attention distribution exhibits a striking sparsity, where the vast majority of genes contribute minimally while a small fraction of “anchor” genes receive weights significantly exceeding the 0.02 threshold. This selective focus demonstrates the model’s capacity to identify and leverage key genetic determinants for metabolic similarity prediction, rather than treating the genome as a simple collection of genes. Notably, the two highly attended genes, SAMN05660413 03200 and SAMN05660413 01847, are both identified as response regulators of two-component systems. These systems serve as essential “environmental sensors” that allow bacteria to coordinate their internal metabolic flux in response to external stimuli. By assigning peak attention to these master regulatory elements, MicroGenomer demonstrates an advanced ability to prioritize the genomic control hubs that govern metabolic similarity, effectively filtering out housekeeping genetic background to focus on the drivers of functional divergence.

To characterize the structure of the gene-level embedding space, we aggregate gene embeddings from all 5,587 bacterial genome-scale metabolic models and cluster them into 512 distinct groups. We then focus on the top 50 clusters, ranked by the number of member genes, and randomly sample 50 gene embeddings from each for visualization. As shown in Fig. 4C, the sampled gene embeddings (represented by dots) and their corresponding cluster centers (represented by stars) are projected into a two-dimensional space using t-SNE [49], color-coded by cluster membership. Gene embeddings within the same cluster form compact, well-defined neighborhoods around their respective centers. This spatial organization demonstrates that MicroGenomer effectively structures the genomic functional space, grouping genes with similar representational patterns into coherent clusters. Such high-fidelity clustering indicates that the learned embeddings are robustly correlated with functional identity, providing a foundation for the model’s ability to interpret complex metabolic relationships.

We aggregate gene attention weights at the cluster level to identify functional groups that drive the model’s predictions. As shown in Fig. 4D, the mean attention weights for the top 20 clusters reveal that Cluster 27 receives the highest focus from the model. This concentration of attention suggests that Cluster 27 represents a critical functional module that MicroGenomer prioritizes when evaluating metabolic similarity.

To provide a concrete biological interpretation of the high-attention signals, we perform an in-depth functional analysis of Cluster 27, the most influential gene group identified by the attention mechanism. Cross-referencing Cluster 27 with our metabolic model dataset reveals that this module is present in 898 out of 5,587 genomes, with the majority harboring a concentrated set of 2 to 4 genes (Fig. 4E). We characterize this cluster by performing functional annotation with Prokka and metabolic pathway reconstruction using KofamScan and KEGG Mapper. Quantitative analysis of 1,193 CDSs within this cluster demonstrates a high density of metabolism-related elements, where 38.7% of these genes are either directly assigned to metabolic COG classes or carry specific Enzyme Commission (EC) numbers. These genes cover essential metabolic modules, including carbon metabolism (e.g., atoC and fadR), nitrogen assimilation (ntrC/glnG), and phosphate starvation response (phoP). KEGG reconstruction further supports this metabolic focus, consistently recovering global “Metabolic pathways” (map01100) and “Purine metabolism” (map00230), alongside 14 distinct two-component systems (map02020) that act as signal transduction interfaces to reroute metabolic flux in response to environmental cues. Taken together, these findings confirm that the high-attention focus of MicroGenomer is not distributed randomly but is precisely anchored to a conserved metabolic regulatory network, capturing the core functional hubs that govern nutrient utilization and environmental adaptation.

### 2.5 MicroGenomer facilitates experimentally validated discoveries of microbial traits

To evaluate the predictive power of MicroGenomer in real-world scenarios, we utilize the model to perform trait predictions across a candidate reservoir of 23 newly sequenced isolates [50]. We focus our experimental validation on optimal growth temperature and pH, as these represent the most fundamental environmental determinants for microbial cultivation. To ensure the validity of these predictions, we first establish a baseline by aligning our laboratory cultivation conditions with the metadata of representative species from the training set. As shown in Fig. 5A and Fig. 5C, the measured growth optima for these reference strains, which span three distinct phyla and four distinct genera, precisely match their database labels, confirming that our cultivation system successfully replicates the environmental context modeled by MicroGenomer.

**Fig. 5:**
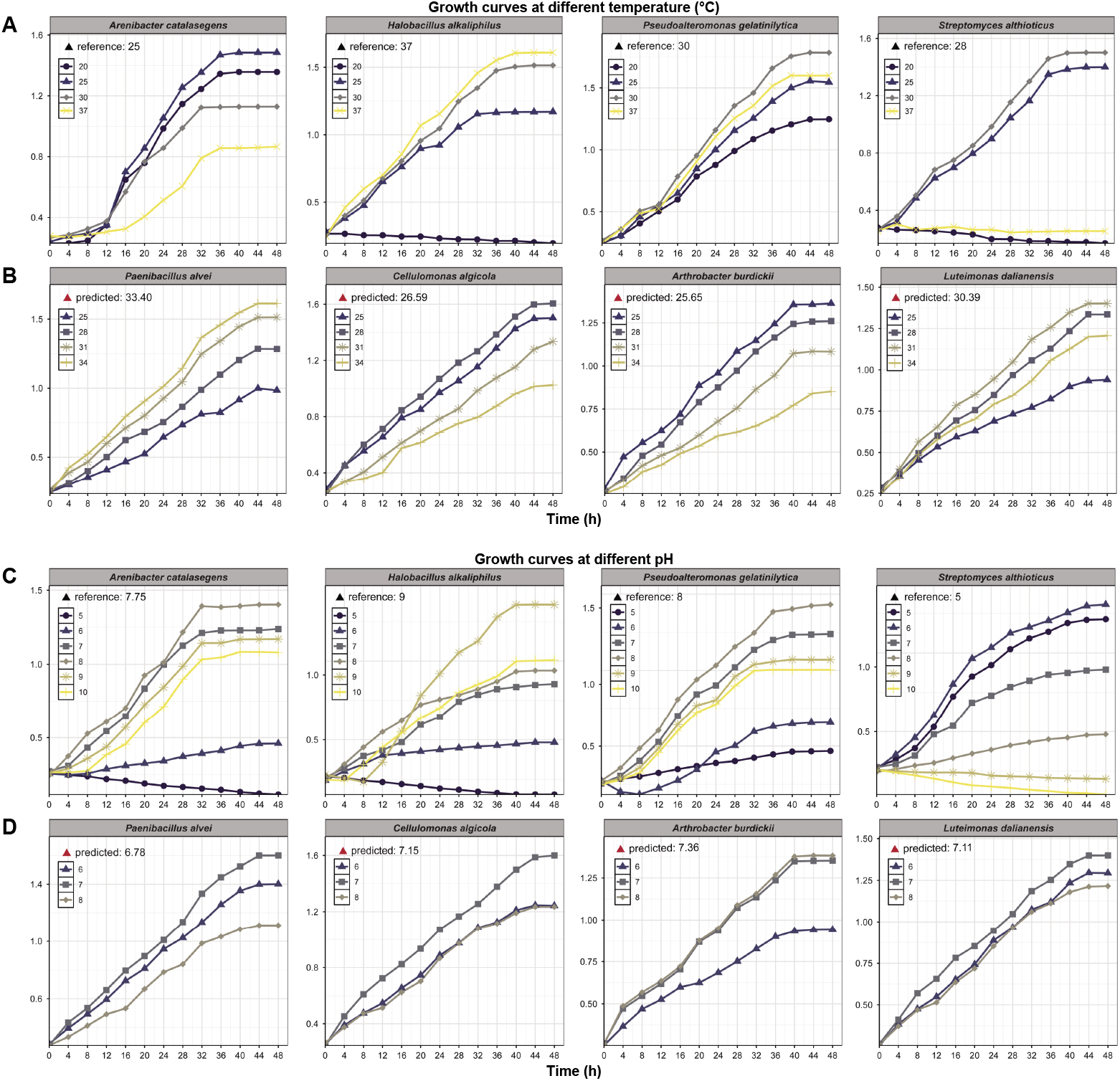
Wet-lab validation for predicted ecophysiological traits of MicroGenomer. **(A)** Calibration of temperature cultivation system using reference strains. **(B)** MicroGenomer temperature predictions vs. experimental growth profiles. **(C)** Calibration of pH cultivation system using reference strains. **(D)** MicroGenomer pH predictions vs. experimental growth profiles.

With the system calibrated, we perform targeted validation on four representative strains selected from the candidate reservoir based on their taxonomic correspondence to the benchmark species. The experimental results reveal that the optimal temperature (Fig. 5B) and pH (Fig. 5D) for these previously uncharacterized isolates are in high concordance with the model’s trait predictions. This consistent accuracy across diverse taxa demonstrates that MicroGenomer can effectively translate raw genomic sequences into actionable physiological insights. By significantly narrowing the experimental search space, the model provides a powerful tool for the directed screening and functional discovery of novel microbial lineages.

## 3 Discussion

The vast diversity of microbial life represents an untapped frontier for biological discovery and a cornerstone of the global biomanufacturing revolution. However, deciphering the complex mapping from massive, fragmented genetic sequences to macroscopic ecophysiological traits remains a fundamental challenge. In this work, we present MicroGenomer, a foundation model that establishes a new paradigm for genomic research by bridging the gap from gene-scale to genome-scale and ultimately to organismal phenotypes. By distilling information from extensive genomic corpora into unified, high-dimensional representations, MicroGenomer moves beyond traditional single-marker analysis to provide a holistic view of a microbe’s functional potential. Instead of relying on isolated genetic features, our approach redefines the representation of life at the genomic level by integrating the collective influence of the coding space. This whole-genome representation allows us to distinguish evolutionary similarities and functional divergences across diverse taxa with unprecedented resolution, offering a robust framework for streamlining strain screening and optimizing cultivation strategies in biotechnology.

MicroGenomer’s core contribution lies in its ability to aggregate thousands of discrete genetic signals into a unified genome-scale embedding, providing a novel lens through which to predict complex ecophysiological traits. Traditional methods often rely on predefined feature engineering or localized genetic information. In contrast, by integrating the entire coding space, MicroGenomer captures global patterns governing strain attributes such as oxygen tolerance, optimal temperature, and salinity. This leap from gene-centric to genome-centric modeling enables the model to understand how functional evolution varies across different lineages. Even under conditions of severe label scarcity, such as optimal pH prediction, this representation, which fuses evolutionary breadth with functional depth, demonstrates robustness superior to traditional feature-rich baselines. This suggests that genome-scale representations can uncover universal biological laws that govern microbial existence, transcending the limitations of classical phylogenetic methods.

Through attention mechanisms and clustering analysis in the representation space, we are able to re-examine the functional trajectories of microbial evolution. MicroGenomer is not a “black-box” predictor, the structural organization of its embedding space aligns closely with taxonomic branching and metabolic similarity. This consistency reveals a profound biological phenomenon: the model spontaneously anchors its attention on “functional hubs”, highly conserved and central modules, such as the signal transduction and energy metabolism elements found in Cluster 27. This capability provides a new research paradigm for genomics, allowing researchers to identify key genetic determinants of ecological adaptation and distinguish evolutionary functional differences even within uncharacterized clades. This sequence-based, phylogeny-aware representation learning offers a transparent analytical solution for deconstructing complex genotype-phenotype associations.

Despite these advancements, mapping sequence patterns to complex functional logic remains an ongoing challenge. Current DNA language models often operate as sequence-level encoders focused on nucleotide patterns. While MicroGenomer transcends this by implementing a gene-centric aggregation mechanism, explicit modeling of non-coding regulatory elements and gene synteny remains an area for future exploration. Furthermore, the taxonomic bias and sparsity of available phenotype data continue to limit generalization across rare evolutionary branches. Future extensions will aim to integrate operon structures, multi-omics data, and uncertainty estimation into a unified framework. As more diverse genomes with precise phenotypic annotations accumulate, this lightweight and adaptable foundation model will serve as an extensible biological resource. It is poised to integrate deeply into laboratory and industrial pipelines, driving the transition of microbiology from a descriptive science to a predictive and creative discipline.

## Data and code availability

The OpenGenome dataset [51], the GTDB dataset [4, 52], the bac120 phylogenetic tree [34], and all datasets used for downstream tasks are available from the corresponding original studies. Microbial ecophysiological trait data were obtained from GenomeSPOT, Phydon, gRodon, and iProbiotics, all of which are publicly accessible. The GenomeSPOT software package is available at https://github.com/cultivarium/GenomeSPOT. The Phydon package can be accessed at https://github.com/xl0418/ Phydon, with analysis code archived at https://github.com/xl0418/PhydonAnalysis. The gRodon package, including documentation and a vignette, is available at https://github.com/jlw-ecoevo/gRodon. The iProbiotics database can be accessed at http://bioinfor.imu.edu.cn/iprobiotics.

## Acknowledgements

This work was supported by the National Key R&D Program of China (No. 2024YFA0919700), the National Natural Science Foundation of China (No. 32400456), and the National Key R&D Program of China (No.2024YFC3505100).

## 4 Methods

### 4.1 Datasets and data processing

For large-scale pre-training, we utilize the OpenGenome corpus, comprising 234.5 billion nucleotides, to capture the fundamental diversity of microbial genomic sequences. To bridge the gap between raw sequences and functional phenotypes, we implement a microorganism-focused mid-training stage using a curated GTDB microbial marker gene collection. This dataset contains 36 billion nucleotides of CDSs from approximately 110k dereplicated genomes, covering 53 archaeal and 120 bacterial marker gene families. By focusing on these conserved coding elements, we steer representation learning toward the genetic determinants most directly linked to metabolic and physiological functions. For post-training and downstream evaluation, we evaluate MicroGenomer across a spectrum of ecophysiological trait prediction tasks derived from three established benchmarks: iProbiotics (probiotic identity) [36], GenomeSPOT (oxygen tolerance, optimal temperature, salinity, and pH) [37], and Phydon (maximum growth rate) [38]. These tasks involve a range of labeled genomes (405–3,637), reflecting the typical scarcity of experimentally characterized traits in microbiology. Additionally, for wet-lab validation, we obtain genome assemblies for 23 newly collected microbial isolates [50] to verify the model’s practical predictive utility.

For all stages, input sequences are tokenized over a vocabulary of five symbols: the four canonical nucleotides (A, T, C, G) and a special symbol N representing unknown bases. Tokens are then mapped to embedding vectors and fed into the model.

For large-scale DNA pre-training, we adopt an MLM objective. Given a sequence, we randomly select 15% of tokens and apply the standard 80/10/10 replacement strategy: 80% of selected tokens are replaced by a special `<mask>` token, 10% are replaced by a random token from the vocabulary, and the remaining 10% are left unchanged. The model is trained to recover the original nucleotides from their surrounding context. Genomes from the OpenGenome corpus are segmented into fixed-length windows of 8,192 bp. During training, we sample windows by sliding along each genome and trimming at the ends as needed, for held-out evaluation, we use non-overlapping 8,192-bp windows drawn from reserved genomic regions so that pre-training performance is measured on unseen sequence segments.

For microorganism-focused mid-training, we start from the curated GTDB microbial marker gene collection, which contains CDSs from approximately 110,000 dereplicated genomes. We partition genomes into three disjoint sets at the genome level. First, an out-of-distribution (OOD) test set is defined at the taxonomic rank of order: all genomes belonging to 368 orders that are absent from the training set are held out, yielding roughly 5,000 genomes for OOD evaluation. Second, from the remaining genomes, we draw an i.i.d. validation set of about 11,000 genomes, and the rest (about 84,000 genomes) form the mid-training set. Evolutionary distances between genomes are derived from the bac120 reference tree [34], which is constructed by concatenating 120 single-copy marker genes and inferring a maximum-likelihood tree with FastTree [33]. Branch lengths from this tree are used as supervision for the phylogeny-aware loss in the mid-training stage.

For microbial ecophysiological trait prediction, we follow phylogenetically informed splitting protocols that closely mirror those used in the original benchmark studies. For the iProbiotics dataset [36], we first construct a phylogenetically separated test set and then perform five-fold cross-validation on the remaining genomes, ensuring that the independent test set is never mixed with training folds; experiments are run at both the species and genus levels. For the four GenomeSPOT tasks (oxygen tolerance and optimal temperature, salinity, and pH) [37], we adopt the family-level holdout designto simulate prediction for phylogenetically novel taxa, ensuring that evaluation reflects the model’s capacity for cross-clade generalization. For maximum growth rate prediction, we follow the phylogenetically blocked ten-fold cross-validation protocol introduced in Phydon [38], using the authors’ clade-level fold assignments: in each fold, models are trained on genomes from nine phylogenetic blocks and evaluated on the remaining block, and results are aggregated over all ten folds. Together, these partitioning schemes ensure that evaluation reflects both within-clade generalization and transfer to phylogenetically distant taxa.

### 4.2 MicroGenomer architecture and training objectives

MicroGenomer is trained in three stages: a large-scale DNA pre-training stage, a phylogeny-aware mid-training stage, and a task-specific post-training stage. This foundation model is built on a Transformer architecture designed for long DNA sequences and microbial genomes. The model operates on a vocabulary of five symbols (A, T, C, G, and N for unknown bases), which are mapped to token embeddings and processed by a stack of self-attention and feed-forward blocks. The model has approximately 470 million parameters, with 24 Transformer layers, a hidden size of 1280, a feed-forward dimension of 3413, 20 attention heads, and a maximum sequence length of 8192 tokens. The learning rate is set to 4 *×* 10^−4^.

The feed-forward sublayers use the Gated Gaussian Error Linear Unit (GEGLU) activation, a variant of the Gated Linear Unit (GLU) in which the gating mechanism is parameterized by a GELU nonlinearity [53]. This activation improves optimization stability and accuracy in large Transformer-based language models. To further stabilize deep training, we combine Layer Normalization with DeepNorm-style scaling [54], which helps control gradient magnitudes and mitigates vanishing or exploding gradients in deep Transformer stacks. For positional encoding, we employ Rotary Positional Embeddings (RoPE) [55] together with Dynamic NTK scaling, enabling the model to represent long-range dependencies and to generalize smoothly across different sequence lengths.

In the pre-training stage, MicroGenomer is trained on genome windows of length 8,192 using an MLM objective. Given an input sequence **x**, a subset of positions ℳ is selected for prediction, and the model is trained to reconstruct the masked nucleotides from their context. The MLM loss is

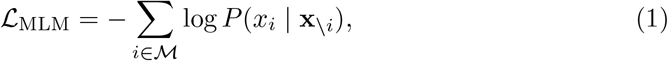

where *x*_*i*_ is the true nucleotide at position *i* and **x\**_*i*_ denotes the sequence with position *i* masked. During this stage, 15% of tokens are selected with the standard 80/10/10 replacement strategy (mask / random token / unchanged), as described in the data processing section.

In the mid-training stage, the model is further adapted on GTDB marker-gene CDSs with supervised objectives that align genome representations with evolutionary distances and gene categories. Let *d*_evo_(*i, j*) denote the evolutionary distance between species *i* and *j*, derived from branch lengths on the bac120 phylogenetic tree [34], and let **z**_*i*_ and **z**_*j*_ be the corresponding species-level embeddings produced by MicroGenomer. We define the representation-space distance as

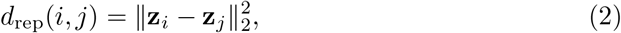

and minimize the discrepancy between representation distances and phylogenetic distances via

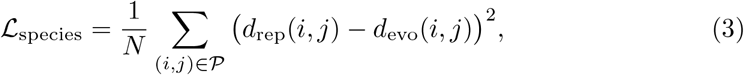

where 𝒫 is the set of sampled species pairs and *N* = |𝒫|. This loss encourages species that are close on the phylogenetic tree to have nearby embeddings, and distant species to be separated in the representation space.

In parallel, we introduce a gene category classification objective over GTDB marker genes. Each gene is assigned to one of 273 marker-gene categories, and a linear probing layer maps its embedding to a categorical distribution. The gene category classification loss is the standard cross-entropy:

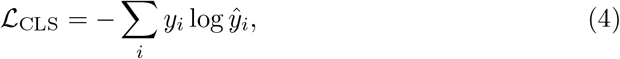

where *y*_*i*_ is the one-hot encoded true label and *ŷ*_*i*_ is the predicted probability for the *i*-th gene. Together, the MLM loss from the pre-training stage and the species-level distance and gene classification losses from the mid-training stage shape a shared genome representation that is sensitive to both local sequence patterns and global evolutionary structure.

In the post-training stage, MicroGenomer is adapted to specific downstream tasks. For sequence-based genomic benchmarks such as the GUE tasks (enhancer–promoter interaction, species classification, and epigenetic mark prediction), we apply parameter-efficient LoRA fine-tuning on top of the pre-trained and mid-trained encoder, and optimize task-specific classification heads using standard cross-entropy losses. For microbial phenotype prediction tasks (e.g., probiotic identity, oxygen tolerance, and optimal temperature, salinity, pH, and growth rate), we treat the MicroGenomer encoder as a frozen feature extractor: genome-level embeddings derived from CDSs are fed into lightweight regression or classification heads (MLPs and classical machine-learning models), which are trained with cross-entropy losses for classification and mean squared error losses for regression. For the metabolic similarity learning task, we attach an attention-pooling layer over gene embeddings and a regression head that predicts pairwise metabolic similarity scores between genomes, optimized with a mean squared error objective with respect to similarities derived from genome-scale metabolic models. These post-training heads allow MicroGenomer to support a range of ecophysiological traits and representation-analysis tasks while reusing a single shared genome encoder.

### 4.3 Downstream task details

#### Zero-shot protein and RNA fitness prediction

Zero-shot prediction enables AI models to perform tasks on categories or data that are not explicitly seen during training by leveraging similarities to previously learned patterns [21]. This is particularly valuable when labeled data are scarce, as models can adapt to new scenarios with minimal or no additional training. For protein DMS, we start from the ProteinGym suite [22] and retain every experiment in which the wild-type codon context can be unambiguously reconstructed. This yields six prokaryotic datasets (*Firnberg, Jacquier, Kelsic, Weeks, Rockah*, and *Chen*) [23]. To enable fair comparison between nucleotide and protein language models, we process both nucleotide- and amino-acid–level DMS data in a consistent way. When the original studies report both nucleotide and protein sequences, we use the nucleotide-based fitness values as ground truth; in cases where discrepancies arise, we prioritize nucleotide measurements and evaluate protein models on the corresponding translated sequences. Mutations introducing stop codons are retained for nucleotide models but excluded from protein-model evaluation to ensure a comparable label space [24, 51].

For ncRNA, we follow the protocol in [51] and curate seven DMS benchmarks (*Zhang, Pitt, Hayden, Guy, Kobori, Domingo*, and *Andreasson*). These datasets provide experimentally measured fitness or activity for large panels of single and, in some cases, combinatorial mutations, enabling a systematic assessment of zero-shot mutational effect prediction on non-coding RNAs. Dataset sizes are summarized in Tab. 1.

**Table 1:**
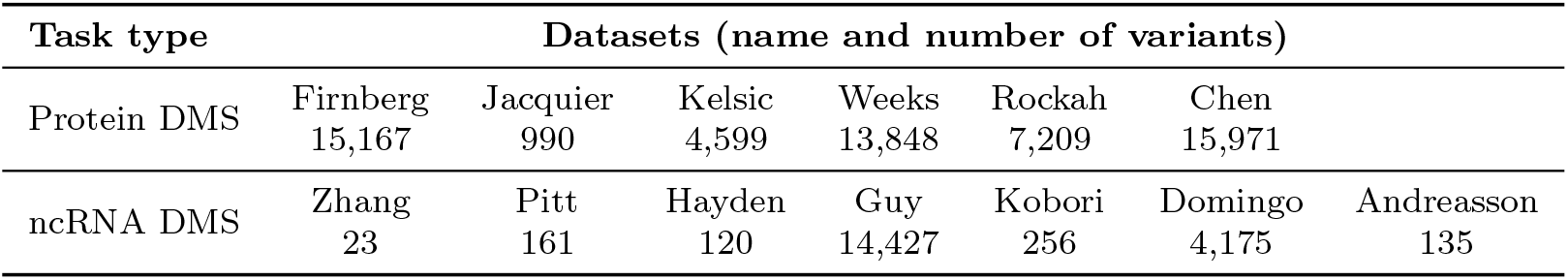
Protein and ncRNA DMS dataset statistics.

| Task type | Datasets (name and number of variants) |  |  |  |  |  |  |
| --- | --- | --- | --- | --- | --- | --- | --- |
| Protein DMS | Firnberg<br>15,167 | Jacquier<br>990 | Kelsic<br>4,599 | Weeks<br>13,848 | Rockah<br>7,209 | Chen<br>15,971 |  |
| ncRNA DMS | Zhang<br>23 | Pitt<br>161 | Hayden<br>120 | Guy<br>14,427 | Kobori<br>256 | Domingo<br>4,175 | Andreasson<br>135 |

#### GUE benchmarks

We evaluate MicroGenomer on the GUE benchmarks [23], which includes epigenetic mark prediction, enhancer–promoter interaction, and species classification tasks. For epigenetic mark prediction in yeast, we use ten datasets and split each into training, validation, and test sets in an 8:1:1 ratio. For enhancer–promoter interaction, the input consists of paired enhancer and promoter sequences from the human genome, and the model predicts a binary interaction label. For species classification, we construct datasets from viral and fungal reference genomes downloaded from GenBank [56], and train MicroGenomer to assign each genomic segment to its species label.

On all GUE tasks, we fine-tune MicroGenomer with parameter-efficient LoRA [28]. Unless otherwise specified, we use a learning rate of 1 × 10^−3^ and a batch size of 32. LoRA rank and scaling factor are set to LoRA-*r* = 4 and LoRA-*α* = 32, respectively. We insert LoRA adapters into the two MLP layers of the task-specific classification head (hidden dimensions 256 and 128), while keeping the shared MicroGenomer encoder fixed.

#### Microbial phenotype prediction tasks

For microbial phenotype prediction, we use three groups of tasks derived from published benchmarks: probiotic identification, environmental growth-condition prediction, and maximum growth rate estimation.

For probiotic identification, we use the iProbiotics dataset [36], which provides species- and genus-level labels indicating whether a strain is probiotic. We adopt the phylogenetically informed splitting strategy, performing experiments at both the species and genus levels and comparing MicroGenomer with iProbiotics and MLC [39]. Prediction heads are trained with cross-entropy loss.

For oxygen tolerance and optimal temperature, salinity, and pH preference, we use the four GenomeSPOT tasks [37]. We follow the GenomeSPOT protocol and retain a family-level held-out test set to evaluate out-of-distribution generalization. The remaining genomes are split into training and validation sets, and experiments are conducted at the species level. MicroGenomer provides frozen genome embeddings, which are fed into lightweight regression or classification heads trained with mean squared error (for continuous traits) or cross-entropy loss (for categorical traits). GenomeSPOT serves as the primary baseline.

For maximum growth rate prediction, we use the dataset and phylogenetically blocked ten-fold cross-validation scheme introduced in Phydon [38]. We directly adopt the original clade-based fold assignments, train prediction heads on nine folds, and evaluate on the held-out fold. Performance is reported using RMSE, MAE, and Spearman correlation, and MicroGenomer is compared with Phydon, NNM, gRodon, and phylopred [19, 38]. In all cases, the MicroGenomer encoder remains frozen and only the downstream heads are trained.

### 4.4 Wet-lab validation processing

#### Strains selection

The experimental validation follows a strategic pipeline designed to test both protocol consistency and predictive novelty. We first construct a candidate reservoir by performing whole-genome sequencing and trait predictions on 23 newly isolated strains [50]. We prioritize the validation of temperature and pH optima due to their critical role in defining microbial niches. For protocol alignment, we select four reference species (Arenibacter catalasegens, Halobacillus alkaliphilus, Pseudoalteromonas gelatinilytica, and Streptomyces althioticus) with known labels from the GenomeSPOT dataset [37]. These serve as positive controls to calibrate the laboratory environment. For independent validation, we then select four representative strains from the 23-isolate reservoir (Paenibacillus alvei, Arthrobacter burdickii, Cellulomonas algicola, and Luteimonas dalianensis). These strains are chosen based on two criteria: (1) taxonomic diversity spanning three distinct phyla and four distinct genera to rigorously evaluate cross-clade generalization, and (2) complete absence from the model’s training and initial testing phases to ensure a strictly unbiased evaluation. All taxonomic identities are independently confirmed via 16S rRNA gene sequence analysis.

#### Cultivation and gradient assay design

Isolates are cultured in 2216E liquid medium, with pH adjusted via 0.1 mol/L HCl/-NaOH and monitored by a calibrated pH meter. To ensure the precision of the optimal condition identification, we design a dense gradient of experimental conditions. For the alignment strains, the gradient is centered on literature-reported values; for the validation reservoir strains, it is centered on the model-derived trait predictions. Assays are performed in 96-well plates using an alternating well pattern (checkerboard layout) to minimize heat and metabolite interference between samples. Each isolate is tested in triplicate across at least five gradient points for both temperature and pH. Growth is quantified by measuring Optical Density (OD) at 600 nm every 4 hours over a 48-hour incubation period. The optimal growth points are determined by comparing the maximum growth rates and stationary phase densities derived from the resulting growth curves.

### 4.5 Evaluation criteria

For zero-shot protein and ncRNA fitness prediction tasks, we evaluate models using the Spearman correlation between experimentally measured fitness values and model-derived sequence scores. For autoregressive language models, we compute sequence scores from log-likelihoods under a consistent masking scheme. For masked language models such as MicroGenomer, we derive pseudo-likelihood scores by comparing the model’s token-level probabilities at mutated positions (equivalently, the perplexity difference between wild-type and mutant sequences restricted to the mutated sites). In all cases, performance is summarized by the Spearman correlation between these scores and experimental fitness measurements.

For regression tasks (e.g., optimal temperature, salinity, pH, and maximum growth rate), we compare predicted values *ŷ*_*i*_ with ground-truth measurements *y*_*i*_ using RMSE, MAE, and Spearman correlation. Let *N* be the number of samples. RMSE and MAE are defined as

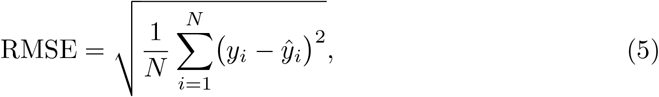

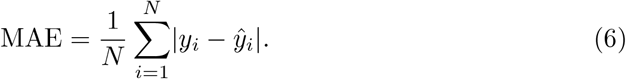

Spearman correlation is computed on the rank-transformed values *r*(*y*_*i*_) and *r*(*ŷ*_*i*_),

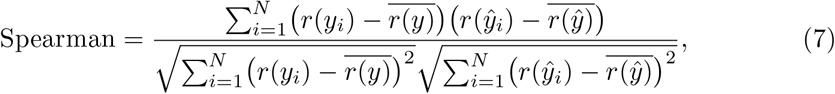

where *r*(*y*_*i*_) and *r*(*ŷ*_*i*_) denote the ranks of *y*_*i*_ and *ŷ*_*i*_, and 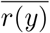 and 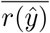 are the mean ranks.

For classification tasks (e.g., probiotic identification, oxygen tolerance, and enhancer–promoter interaction), we use the Matthews Correlation Coefficient (MCC), F1 score, and Area Under the ROC Curve (AUC). Given true positives (*TP*), true negatives (*TN*), false positives (*FP*), and false negatives (*FN*), MCC is defined as

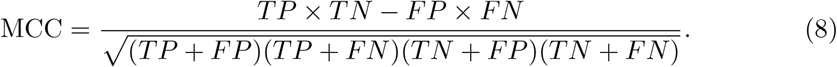

The F1 score is the harmonic mean of precision and recall,

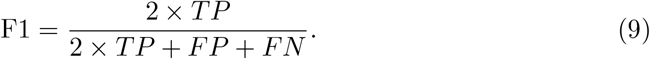

AUC measures the probability that a randomly chosen positive sample receives a higher prediction score than a randomly chosen negative sample, summarizing performance across all classification thresholds.

For clustering-based evaluations (e.g., genus-level clustering of genome embeddings), we report the Adjusted Rand Index (ARI), which measures the similarity between predicted clustering assignments and ground-truth labels, adjusted for chance. A higher ARI indicates better alignment,

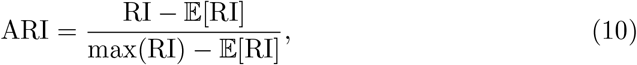

where RI is the Rand Index, defined based on pairwise agreements and disagreements between two partitions, 𝔼 [RI] is its expected value under random label assignments.

### 4.6 Project implementation

Our experiments are conducted on a computing cluster with 8 host machines, each equipped with 8 NVIDIA A100 GPUs, providing powerful parallel processing. The cluster also has 128 CPU cores and 1007 GB of system memory. All machines run Ubuntu with kernel version 5.4.0-139-generic. We use RDMA (Remote Direct Memory Access) for optimized inter-node communication. The GPU driver version is DCGM 525.147.05. Our training pipeline is built on top of the Megatron and DeepSpeed frameworks, integrating FlashAttention to accelerate attention computation. For implementation, we rely on PyTorch and PyTorch Lightning, both of which offer flexibility and robustness for large-scale deep learning research. For the training settings, we set ZERO STAGE to 1 to reduce memory usage by partitioning the optimizer states. To accelerate the training process, we use the Brain Floating Point 16 (BF16) data type for the model parameters. Meanwhile, we keep the gradient accumulation data type as FP32 to ensure numerical stability.

## Notes

### Competing Interest Statement

The authors have declared no competing interest.

## References

[1] Crowther, T.W., Rappuoli, R., Corinaldesi, C., Danovaro, R., Donohue, T.J., Huisman, J., Stein, L.Y., Timmis, J.K., Timmis, K., Anderson, M.Z., Bakken, L.R., Baylis, M., Behrenfeld, M.J., Boyd, P.W., Brettell, I., Cavicchioli, R., Delavaux, C.S., Foreman, C.M., Jansson, J.K., Koskella, B.: Scientists’ call to action: Microbes, planetary health, and the sustainable development goals. Cell 187(19), 5195–5216 (2024) 10.1016/j.cell.2024.07.051

[2] Thompson, L.R., Sanders, J.G., McDonald, D., Amir, A., Ladau, J., Locey, K.J., Prill, R.J., Tripathi, A., Gibbons, S.M., Ackermann, G., Navas-Molina, J.A., Janssen, S., Kopylova, E., Vázquez-Baeza, Y., González, A., Morton, J.T., Mirarab, S., Xu, Z.Z., Jiang, L., Haroon, M.F., Kanbar, J., Zhu, Q., Song, S.J., Kosciolek, T., Bokulich, N.A., Lefler, J., Brislawn, C.J., Humphrey, G., Owens, S.M., Hampton-Marcell, J., Berg-Lyons, D., McKenzie, V., Fierer, N., Fuhrman, J.A., Clauset, A., Stevens, R.L., Shade, A., Pollard, K.S., Goodwin, K.D., Jansson, J.K., Gilbert, J.A., Knight, R., Consortium, T.E.M.P.: A communal catalogue reveals earth’s multiscale microbial diversity. Nature 551, 457–463 (2017) 10.1038/nature24621

[3] Nayfach, S., Roux, S., Seshadri, R., Udwary, D., Varghese, N., Schulz, F., Wu, D., Paez-Espino, D., Chen, I.-M., Huntemann, M., Palaniappan, K., Ladau, J., Mukherjee, S., Reddy, T.B.K., Nielsen, T., Kirton, E., Faria, J.P., Edirisinghe, J.N., Henry, C.S., Jungbluth, S.P., Chivian, D., Dehal, P., Wood-Charlson, E.M., Arkin, A.P., Tringe, S.G., Visel, A., Consortium, I.D., Woyke, T., Mouncey, N.J., Ivanova, N.N., Kyrpides, N.C., Eloe-Fadrosh, E.A.: A genomic catalog of earth’s microbiomes. Nature Biotechnology 39, 499–509 (2021) 10.1038/s41587-020-0718-6

[4] Parks, D.H., Chuvochina, M., Rinke, C., Mussig, A.J., Chaumeil, P.-A., Hugenholtz, P.: Gtdb: an ongoing census of bacterial and archaeal diversity through a phylogenetically consistent, rank normalized and complete genome-based taxonomy. Nucleic Acids Research 50(D1), 785–794 (2022) 10.1093/nar/gkab776

[5] Richardson, L., Allen, B., Baldi, G., Beracochea, M., Bileschi, M.L., Burdett, T., Burgin, J., Caballero-Pérez, J., Cochrane, G., Colwell, L.J., Curtis, T., Escobar-Zepeda, A., Gurbich, T.A., Kale, V., Korobeynikov, A., Raj, S., Rogers, A.B., Sakharova, E., Sanchez, S., Wilkinson, D.J., Finn, R.D.: Mgnify: the microbiome sequence data analysis resource in 2023. Nucleic Acids Research 51(D1), 753–759 (2023) 10.1093/nar/gkac1080

[6] Cébron, A., Zeghal, E., Usseglio-Polatera, P., Meyer, A., Bauda, P., Lemmel, F., Leyval, C., Maunoury-Danger, F.: Bactotraits – a functional trait database to evaluate how natural and man-induced changes influence the assembly of bacterial communities. Ecological Indicators 130, 108047 (2021) 10.1016/j.ecolind.2021.108047

[7] Latif, A., zad, A.S., Niazi, S., Zahid, A., Ashraf, W., Iqbal, M.W., Rehman, A., Riaz, T., Aadil, R.M., Khan, I.M., Özogul, F., Rocha, J.M., Esatbeyoglu, T., Korma, S.A.: Probiotics: mechanism of action, health benefits and their application in food industries. Frontiers in Microbiology 14, 1216674 (2023) 10.3389/fmicb.2023.1216674

[8] Pandey, R.P., Gunjan, Himanshu, Mukherjee, R., Chang, C.-M.: Nanocarrier-mediated probiotic delivery: a systematic meta-analysis assessing the biological effects. Scientific Reports 14, 631 (2024)

[9] Orouji, S., Liu, M.C., Korem, T., Peters, M.A.K.: Domain adaptation in small-scale and heterogeneous biological datasets. Science Advances 10(51), 6040 (2024) 10.1126/sciadv.adp6040

[10] Jumper, J., Evans, R., Pritzel, A., Green, T., Figurnov, M., Ronneberger, O., Tunyasuvunakool, K., Bates, R., Žídek, A., Potapenko, A., Bridgland, A., Meyer, C., Kohl, S.A.A., Ballard, A.J., Cowie, A., Romera-Paredes, B., Nikolov, S., Jain, R., Adler, J., Back, T., Petersen, S., Reiman, D., Clancy, E., Zielinski, M., Steinegger, M., Pacholska, M., Berghammer, T., Bodenstein, S., Silver, D., Vinyals, O., Senior, A.W., Kavukcuoglu, K., Kohli, P., Hassabis, D.: Highly accurate protein structure prediction with alphafold. Nature 596, 583–589 (2021) 10.1038/s41586-021-03819-2

[11] Baek, M., DiMaio, F., Anishchenko, I., Dauparas, J., Ovchinnikov, S., Lee, G.R., Wang, J., Cong, Q., Kinch, L.N., Schaeffer, R.D., Millán, C., Park, H., Adams, C., Glassman, C.R., DeGiovanni, A., Pereira, J.H., Rodrigues, A.V., Dijk, A.A., Ebrecht, A.C., Opperman, D.J., Sagmeister, T., Buhlheller, C., Pavkov-Keller, T., Rathinaswamy, M.K., Dalwadi, U., Yip, C.K., Burke, J.E., Garcia, K.C., Grishin, N.V., Adams, P.D., Read, R.J., Baker, D.: Accurate prediction of protein structures and interactions using a three-track neural network. Science 373(6557), 871–876 (2021) 10.1126/science.abj8754

[12] Lin, Z., Akin, H., Rao, R., Hie, B., Zhu, Z., Lu, W., Smetanin, N., Verkuil, R., Kabeli, O., Shmueli, Y., Santos Costa, A., Fazel-Zarandi, M., Sercu, T., Candido, S., Rives, A.: Evolutionary-scale prediction of atomic-level protein structure with a language model. Science 379(6637), 1123–1130 (2023) 10.1126/science.ade2574

[13] Cui, H., Wang, C., Maan, H., Pang, K., Luo, F., Duan, N., Wang, B.: scgpt: toward building a foundation model for single-cell multi-omics using generative ai. Nature Methods 21, 1470–1480 (2024) 10.1038/s41592-024-02201-0

[14] Ji, Y., Zhou, Z., Liu, H., Davuluri, R.V.: Dnabert: pre-trained bidirectional encoder representations from transformers model for dna-language in genome. Bioinformatics 37(15), 2112–2120 (2021) 10.1093/bioinformatics/btab083

[15] Dalla-Torre, H., Gonzalez, L., Mendoza-Revilla, J., Carranza, N.L., Grzywaczewski, A.H., Oteri, F., Dallago, C., Trop, E., Almeida, B.P., Sirelkhatim, H., Richard, G., Skwark, M., Beguir, K., Lopez, M., Pierrot, T.: Nucleotide transformer: building and evaluating robust foundation models for human genomics. Nature Methods 22, 287–297 (2025) 10.1038/s41592-024-02523-z

[16] Nguyen, E., Poli, M., Faizi, M., Thomas, A., Wornow, M., Birch-Sykes, C., Massaroli, S., Patel, A., Rabideau, C., Bengio, Y., Ermon, S., Ré, C., Baccus, S.: Hyenadna: Long-range genomic sequence modeling at single nucleotide resolution. In: Advances in Neural Information Processing Systems, vol. 36, pp. 43177–43201. Curran Associates, Inc., New Orleans (2023)

[17] Hwang, Y., Cornman, A.L., Kellogg, E.H., Ovchinnikov, S., Girguis, P.R.: Genomic language model predicts protein co-regulation and function. Nature Communications 15, 2880 (2024) 10.1038/s41467-024-46947-9

[18] Weimann, A., Mooren, K., Frank, J., Pope, P.B., Bremges, A., McHardy, A.C.: From genomes to phenotypes: Traitar, the microbial trait analyzer. mSystems 1(6), 10–11280010116 (2016) 10.1128/msystems.00101-16

[19] Weissman, J.L., Shengwei Hou, J.A.F.: Estimating maximal microbial growth rates from cultures, metagenomes, and single cells via codon usage patterns. Proceedings of the National Academy of Sciences 118(12), 2016810118 (2021) 10.1073/pnas.2016810118

[20] Diaz, D.J., Kulikova, A.V., Ellington, A.D., Wilke, C.O.: Using machine learning to predict the effects and consequences of mutations in proteins. Current opinion in structural biology 78, 102518 (2023)

[21] Meier, J., Rao, R., Verkuil, R., Liu, J., Sercu, T., Rives, A.: Language models enable zero-shot prediction of the effects of mutations on protein function. Advances in neural information processing systems 34, 29287–29303 (2021)

[22] Notin, P., Kollasch, A., Ritter, D., Van Niekerk, L., Paul, S., Spinner, H., Rollins, N., Shaw, A., Orenbuch, R., Weitzman, R., et al.: Proteingym: Large-scale benchmarks for protein fitness prediction and design. Advances in Neural Information Processing Systems 36, 64331–64379 (2023)

[23] Yang, Q., Guo, Y., Liu, Z., Yang, Y., Yin, Q., Li, S., Ji, S., Chao, L., Zhang, X., Li, S.Z.: Trinitydna: A bio-inspired foundational model for efficient long-sequence dna modeling. arXiv preprint 2507.19229 (2025)

[24] Brixi, G., Durrant, M.G., Ku, J., Poli, M., Brockman, G., Chang, D., Gonzalez, G.A., King, S.H., Li, D.B., Merchant, A.T., et al.: Genome modeling and design across all domains of life with evo 2. BioRxiv, 2025–02 (2025)

[25] Schiff, Y., Kao, C.-H., Gokaslan, A., Dao, T., Gu, A., Kuleshov, V.: Caduceus: Bidirectional equivariant long-range dna sequence modeling. Proceedings of machine learning research 235, 43632–43648 (2024)

[26] Zhou, Z., Ji, Y., Li, W., Dutta, P., Davuluri, R., Liu, H.: Dnabert-2: Efficient foundation model and benchmark for multi-species genome. arXiv preprint 2306.15006 (2023)

[27] Dalla-Torre, H., Gonzalez, L., Mendoza-Revilla, J., Lopez Carranza, N., Grzywaczewski, A.H., Oteri, F., Dallago, C., Trop, E., Almeida, B.P., Sirelkhatim, H., et al.: Nucleotide transformer: building and evaluating robust foundation models for human genomics. Nature Methods, 1–11 (2024)

[28] Hu, E.J., Shen, Y., Wallis, P., Allen-Zhu, Z., Li, Y., Wang, S., Wang, L., Chen, W.: LoRA: Low-Rank Adaptation of Large Language Models (2021). https://arxiv.org/abs/2106.09685

[29] Ji, Y., Zhou, Z., Liu, H., Davuluri, R.V.: Dnabert: pre-trained bidirectional encoder representations from transformers model for dna-language in genome. Bioinformatics 37(15), 2112–2120 (2021)

[30] Machado, D., Andrejev, S., Tramontano, M., Patil, K.R.: Fast automated reconstruction of genome-scale metabolic models for microbial species and communities. Nucleic acids research 46(15), 7542–7553 (2018)

[31] Salahshouri, P., Emadi-Baygi, M., Jalili, M., Khan, F.M., Wolkenhauer, O., Salehzadeh-Yazdi, A.: A metabolic model of intestinal secretions: the link between human microbiota and colorectal cancer progression. Metabolites 11(7), 456 (2021)

[32] Van Oven, M.: Phylotree build 17: Growing the human mitochondrial dna tree. Forensic Science International: Genetics Supplement Series 5, 392–394 (2015)

[33] Price, M.N., Dehal, P.S., Arkin, A.P.: Fasttree: computing large minimum evolution trees with profiles instead of a distance matrix. Molecular biology and evolution 26(7), 1641–1650 (2009)

[34] Parks, D.H., Chuvochina, M., Waite, D.W., Rinke, C., Skarshewski, A., Chaumeil, P.-A., Hugenholtz, P.: A standardized bacterial taxonomy based on genome phylogeny substantially revises the tree of life. Nature biotechnology 36(10), 996–1004 (2018)

[35] Bonnet, E., Peer, Y.: zt: A sofware tool for simple and partial mantel tests. Journal of Statistical software 7, 1–12 (2002)

[36] Sun, Y., Li, H., Zheng, L., Li, J., Hong, Y., Liang, P., Kwok, L.-Y., Zuo, Y., Zhang, W., Zhang, H.: iprobiotics: a machine learning platform for rapid identification of probiotic properties from whole-genome primary sequences. Briefings in Bioinformatics 23(1), 477 (2022)

[37] Barnum, T.P., Crits-Christoph, A., Molla, M., Carini, P., Lee, H.H., Ostrov, N.: Predicting microbial growth conditions from amino acid composition. bioRxiv, 2024–03 (2024)

[38] Xu, L., Zakem, E., Weissman, J.: Improved maximum growth rate prediction from microbial genomes by integrating phylogenetic information. Nature Communications 16(1), 4256 (2025)

[39] Bobbo, T., Biscarini, F., Yaddehige, S.K., Alberghini, L., Rigoni, D., Bianchi, N., Taccioli, C.: Machine learning classification of archaea and bacteria identifies novel predictive genomic features. BMC Genomics 25, 955 (2024) 10.1186/s12864-024-10832-y

[40] Gardner, M.W., Dorling, S.R.: Artificial neural networks (the multilayer perceptron)—a review of applications in the atmospheric sciences. Atmospheric environment 32(14-15), 2627–2636 (1998)

[41] Tibshirani, R.: Regression shrinkage and selection via the lasso. Journal of the Royal Statistical Society Series B: Statistical Methodology 58(1), 267–288 (1996)

[42] McDonald, G.C.: Ridge regression. Wiley Interdisciplinary Reviews: Computational Statistics 1(1), 93–100 (2009)

[43] Drucker, H., Burges, C.J., Kaufman, L., Smola, A., Vapnik, V.: Support vector regression machines. Advances in neural information processing systems 9 (1996)

[44] Breiman, L.: Random forests. Machine learning 45(1), 5–32 (2001)

[45] Cover, T., Hart, P.: Nearest neighbor pattern classification. IEEE transactions on information theory 13(1), 21–27 (1967)

[46] Chen, T., Guestrin, C.: Xgboost: A scalable tree boosting system. In: Proceedings of the 22nd Acm Sigkdd International Conference on Knowledge Discovery and Data Mining, pp. 785–794 (2016)

[47] Cortes, C., Vapnik, V.: Support-vector networks. Machine learning 20(3), 273–297 (1995)

[48] Bergstra, J., Bengio, Y.: Random search for hyper-parameter optimization. The journal of machine learning research 13(1), 281–305 (2012)

[49] Maaten, L.v.d., Hinton, G.: Visualizing data using t-sne. Journal of machine learning research 9(Nov), 2579–2605 (2008)

[50] Dai, M., Zhao, F., Shi, X., Tian, C., Lin, Y., Bai, L., Li, T., Jin, X., Xiao, L., Kristiansen, K., Li, X., Zhang, Z.: Cultivation and sequencing of microbiota members unveil the functional potential of yak gut microbiota. mSystems 10(9), 00367–25 (2025) 10.1128/msystems.00367-25

[51] Nguyen, E., Poli, M., Durrant, M.G., Kang, B., Katrekar, D., Li, D.B., Bartie, L.J., Thomas, A.W., King, S.H., Brixi, G., Sullivan, J., Ng, M.Y., Lewis, A., Lou, A., Ermon, S., Baccus, S.A., Hernandez-Boussard, T., Ré, C., Hsu, P.D., Hie, B.L.: Sequence modeling and design from molecular to genome scale with evo. Science 386(6723), 9336 (2024) 10.1126/science.ado9336

[52] Chaumeil, P.-A., Mussig, A.J., Hugenholtz, P., Parks, D.H.: GTDB-Tk: a toolkit to classify genomes with the Genome Taxonomy Database. Oxford University Press (2020)

[53] Hendrycks, D., Gimpel, K.: Gaussian error linear units (gelus). arXiv preprint 1606.08415 (2016)

[54] Wang, H., Ma, S., Dong, L., Huang, S., Zhang, D., Wei, F.: Deepnet: Scaling transformers to 1,000 layers. IEEE Transactions on Pattern Analysis and Machine Intelligence 46(10), 6761–6774 (2024)

[55] Heo, B., Park, S., Han, D., Yun, S.: Rotary position embedding for vision transformer. In: European Conference on Computer Vision, pp. 289–305 (2024). Springer

[56] Benson, D.A., Cavanaugh, M., Clark, K., Karsch-Mizrachi, I., Lipman, D.J., Ostell, J., Sayers, E.W.: GenBank. Nucleic Acids Research, 41(D1):D36–D42 (2012). 10.1093/nar/gks1195

